# Spinal-level activation of GPR37 in TRPV1-expressing sensory neurons erases nociceptive system sensitization in murine models

**DOI:** 10.1101/2025.11.25.690482

**Authors:** Regan M. Hammond, Jigong Wang, Ramesh Pariyar, Ho Koo, Jun-Ho La

**Author notes:** Both authors contributed equally to this work (Regan M. Hammond, Jigong Wang). Corresponding author: Jun-Ho La, DVM, PhD., Department of Neurobiology, University of Texas Medical Branch, 301 University Blvd., Galveston, TX, USA.

## Abstract

Intense injury induces long-term changes in the spinal nociceptive system, which increases pain in magnitude and duration. We investigated whether activation of G protein-coupled receptor 37 (GPR37) at the spinal level can erase this nociceptive system sensitization and resolve long-lasting, enhanced pain using two animal models: the capsaicin model and the hyperalgesic priming model. Without altering normal mechanical and heat nociception, a single intrathecal (i.th.) administration of two GPR37 agonists, TX14A and protectin D1 (PD1), dose-dependently inhibited capsaicin-induced increase in nociception not only acutely but also long-term. In the hyperalgesic priming model, a single i.th. injection of either GPR37 agonist after an initial injury-induced priming dose-dependently prevented increased nociception following a subsequent inflammatory insult, indicating an unpriming effect. Global GPR37 knockout or conditional knockout of GPR37 in TRPV1-lineage sensory neurons abolished the long-term inhibitory effect and unpriming effect of i.th. TX14A, confirming that GPR37 in this specific cellular population mediates these effects. Ex vivo Ca²⁺ imaging revealed that i.th. TX14A, given in vivo to the capsaicin model a week before imaging, had rescued dorsal horn excitatory and inhibitory interneurons from long-term potentiation and depression of responsiveness to afferent inputs, respectively, suggesting the erasure of capsaicin-induced nociceptive system sensitization. Conditioned place preference tests indicated no obvious abuse liability for centrally administered GPR37 agonists. These findings suggest that spinally targeting GPR37 in TRPV1-expressing sensory neurons represents a promising therapeutic strategy for resolving persistent pain by erasing nociceptive system sensitization.

## Introduction

Chronic pain affects 20% of adults worldwide, with 10% being newly diagnosed with chronic pain each year^16^. This global burden underscores the need for novel therapies that can, for example, resolve pain rather than merely suppressing it temporarily. In this regard, triggering endogenous mechanisms that undo the long-term changes constituting the sensitization of the nociceptive system may represent one potential approach. Proposed here is that G protein-coupled receptor 37 (GPR37) at the spinal level holds the key to this approach, based on the literature presented below.

GPR37 is a parkin-associated endothelin-like receptor predominantly expressed in the central nervous system, particularly in the spinal cord^28^. While GPR37 remains classified as an orphan G protein-coupled receptor, prosaposin and protectin D1 (PD1) have been identified as its putative agonists^22,27,29,45^. Importantly, TX14A, a derivative of prosaposin, has been found to alleviate traumatic, diabetic, chemotherapy-induced, and HIV-associated neuropathic pain-like behaviors in rats^20,21,31^; it is noteworthy that both systemic and intrathecal (i.th.) injections of TX14A were effective, but the former took a long time to reach the maximal effect, compared to the latter, suggesting a delayed access of TX14A (or its metabolite) to its central action site from the periphery. PD1 has also been shown to alleviate various forms of pain-like behaviors via GPR37^4,5,33,42,43^. While the literature mostly focuses on the acute analgesic effects of these GPR37 agonists, it is notable that a single injection of TX14A prevented the development of formalin-induced persistent mechanical hypersensitivity^21^, and the spinal application of PD1 abrogated experimentally induced long-term potentiation (LTP) of C-fiber-evoked field potentials in the dorsal horn^33^, which collectively suggest that GPR37 agonism at the spinal level may have the potential to resolve pain by erasing sensitization of the nociceptive system. In this study, this possibility was tested using two well-characterized animal models of nociceptive system sensitization: the capsaicin model and the hyperalgesic priming model. In rodents, intraplantar (i.pl.) capsaicin injection induces LTP of excitatory postsynaptic responses to high-threshold afferent inputs in the dorsal horn, which does not require ongoing afferent input for maintenance^19^. This long-term change in the dorsal horn aligns with the properties of long-lasting hyperalgesia-like secondary mechanical hypersensitivity in the capsaicin model^25,26,36^. The hyperalgesic priming model manifests nociceptive system sensitization in the form of a primed nociceptive system following an initial acute injury, resulting in increased (in the magnitude and/or duration) nocifensive behaviors upon a subsequent, mild inflammatory insult^23,35^. The priming by an initial injury has also been shown to require molecules needed for long-term synaptic plasticity^11,23^. Using these two animal models and two putative GPR37 agonists, TX14A and PD1, we investigated whether spinal-level GPR37 activation resolves long-lasting, enhanced nociception due to nociceptive system sensitization.

## Methods

### Animals

Adult male and female C57BL/6NCrl mice (aged 7-12 weeks old) were purchased from Charles River (Houston, TX, USA). GPR37 global knockout (KO) mice (JAX# 005806, B6.129P2-*Gpr37*^tm1Dgen^/J), SST^Cre^ mice (JAX# 013044, STOCK *Sst*^tm2.1(cre)Zjh^/J), GAD2^Cre^ mice (JAX# 010802, STOCK *Gad2*^tm2(cre)Zjh^/J), and Ai95D mice (JAX #028865, B6J.Cg-*Gt(ROSA)26Sor*^tm95.1(CAG-GCaMP6f)Hze^/MwarJ) were purchased from Jackson Laboratory and bred in-house. Each Cre mouse line was crossbred with Ai95D mice to produce offspring expressing GCaMP6f in SST- or GAD2-positive neurons for Ca^2+^ imaging. GPR37^fl/fl^ mice (Strain # T005938, C57BL/6JGpt-*Gpr37*^em1Cflox^/Gpt) were purchased from GemPharmatech, and TRPV1^Cre^ (JAX# 017769, B6.129-*Trpv1*^tm1(cre)Bbm^/J) mice were purchased from Jackson Laboratory. The TRPV1^Cre^ mouse line was crossbred with GPR37^fl/fl^ mice to produce F2 offspring (TRPV1^Cre^GPR37^fl/fl^ mice) with GPR37 conditionally knocked out in TRPV1-lineage neurons (i.e., neurons including both transiently and persistently expressing TRPV1). Mice were group-housed (≤5 per cage) under a 12-h light-dark cycle with free access to food and water and randomly assigned to experimental groups. All experimental procedures were approved by the Institutional Animal Care and Use Committee at the University of Texas Medical Branch and in accordance with the National Institutes of Health (NIH) guidelines.

### Animal models of pain

#### Capsaicin model

Under isoflurane anesthesia, a single i.pl. injection of capsaicin (3 μg in 3 μl, in the mixture of 10% ethanol, 10% Tween-20, and 80% saline; Sigma-Aldrich, St. Louis, MO, USA) was given intradermally into the plantar side of one hind paw between the bases of the third and fourth digits using a 30G needle. In mice for ex vivo Ca^2+^ imaging, a piece of gel foam (∼5x10 mm) soaked with either vehicle or 0.1% capsaicin was directly applied to the sciatic nerve under isoflurane anesthesia after skin and muscle incision. Mice were kept anesthetized during the application for 45 minutes, with a sterile gauze covering the incision wound. The incision was then suture-closed, and the spinal cord was harvested for ex vivo Ca^2+^ imaging seven days later.

#### Hyperalgesic priming model

As an initial priming injury, a single dose of interleukin-6 (IL-6, 25 pg in 5 μl, in saline) was intradermally injected into the center of one hind paw using a 30G needle. After IL-6-induced pain hypersensitivity had completely resolved (7 days post-IL-6), a single dose of prostaglandin E_2_ (PGE_2_, 50 ng in 5 μl, in saline) was injected at the same site as IL-6. The ‘not-primed’ group received only a single i.pl. injection of PGE_2_.

Humane endpoints were defined as orbital tightening with failure to move upon stimulation, weight loss ≥20%, or hind limb motor deficit (e.g., foot-dragging).

### Intrathecal drug administration

Mice were anesthetized with 2-2.5% isoflurane, and all intrathecal (i.th.) injections were given between the L5 and L6 vertebrae (lumbar puncture) after removing fur from the injection site using clippers and sterilizing the area using 70% isopropyl alcohol and povidone-iodine wipes. Proper placement of the needle was confirmed by a brief tail-flick response upon insertion. Mice received a single i.th. injection of TX14A (10, 25, or 50 μg in 10 μl, dissolved in saline; cat.# AS-60248-5, Anaspec, Fremont, CA, USA), PD1 (50 or 100 ng in 5 μl, in a mixture of 10% DMSO and 90% saline; item# 10010390, Cayman Chemical, Ann Arbor, MI, USA), or their vehicles 30 min before normal nociception assessment, 30 min after capsaicin injection (in the capsaicin model), or 5 days after IL-6 injection (in the hyperalgesic priming model). In the i.pl. capsaicin model, this time point (30 min post-capsaicin) was chosen to examine how spinal-level GPR37 agonism affects already-sensitized nociceptive system and resultant, ongoing pain hypersensitivity. In the experiments where capsaicin was directly applied to the exposed sciatic nerve to maximize the yield of sensitized dorsal horn neurons from each mouse in ex vivo slice preparations for Ca^2+^ imaging, an additional dose of TX14A was given on the following day (i.e., two injections in total), as this capsaicin application more invasively affected the whole sciatic nerve – potentially producing greater nociceptive system sensitization - than a local i.pl. injection.

In the hyperalgesic priming model, the time point (5 days post-IL-6; when IL-6-induced pain hypersensitivity has resolved) was chosen to avoid interfering with the priming process itself but to specifically examine how spinal-level GPR37 agonism affects the maintenance of a primed state. Experimenters were blinded to the nature of the treatment.

### Behavioral tests

Mice were habituated to the behavioral test conditions and experimenters for 3 days before conducting behavioral procedures.

#### Von Frey filament (vFF) test

Mice were placed in acrylic chambers (14 cm length x 5 cm width x 4.5 cm height) on a raised metal grid-floor platform and were acclimated for 30 min before testing on the day of the experiment. Mechanical sensitivity of the hind paw was tested using 0.98-mN and 9.8-mN vFFs, which evoke approximately 0–20% and 40–60% nocifensive withdrawal responses at baseline in naïve mice, respectively. The mid-hind paw was probed 10 times, and the percentage of withdrawal responses was recorded.

#### Radiant heat (Hargreaves) test

Mice were placed on a glass floor, and a radiant heat source was placed under the glass floor directly under the middle of the afflicted hind paw. The latency (sec) to withdrawal from the heat was automatically detected, and the results from three trials (>1 min intertrial intervals) at a given time-point were averaged.

#### Conditioned place preference (CPP) test

Using a three-chamber apparatus consisting of a bright and dark chamber connected by a smaller grey chamber (place preference LE 892/LE 893, Panlab/Harvard Apparatus, Dover, MA, USA), mice were placed in the small grey chamber and were allowed to freely roam through the apparatus for 30 min. The amount of time that the mouse spent in each chamber was recorded. Next, conditioning was performed once per day for three days. Mice were given i.th. or intracerebroventricular (i.c.v.) injections of TX14A (50 μg), PD1 (100 ng), morphine (7-10 μg), or saline under anesthesia, then placed in their home cage. Once mice fully awoke from anesthesia (15-20 min post-injection), they were placed into the non-preferred (bright) chamber for 30 min. They were also counter-conditioned for 30 min per day by being placed in the dark chamber after waking from anesthesia without receiving any injection on each conditioning day. After the three days of conditioning, mice were placed in the middle grey chamber and allowed to freely roam through the three-chamber apparatus. The change in time (Δt) spent in the drug-paired chamber between the post- and pre-conditioning tests was calculated.

### Western Blot

Proteins were extracted using RIPA buffer (1x RIPA with 100x phosphatase inhibitor and 100x protease inhibitor). Protein concentrations were determined using a BCA assay kit (ThermoScientific, Chicago, IL, USA). Equal quantities of protein were subjected to SDS-PAGE (10% gel) and transferred to a nitrocellulose membrane. Membranes were blocked in 5% skim milk in T-PBS, then incubated overnight at 4°C with antibodies specific to GPR37 (D4C8H, Cell Signaling Technology, Danvers, MA, USA) and GAPDH (D16H11, Cell Signaling Technology). GPR37 antibodies were diluted to 1:1000 with 5% skim milk, GAPDH antibodies were diluted to 1:2000 with Licor blocking buffer (Cat.# 1610734, Bio-Rad, Hercules, CA, USA). Membranes were then incubated with the corresponding secondary antibodies and developed using a chemiluminescent reagent (ThermoScientific).

### Spinal Cord Slice Preparation

Preparation of spinal cord slices with attached dorsal root was done as described previously^32^. Mice were deeply anesthetized using isoflurane, then the lumbar spinal cord was removed and submerged in ice-cold, oxygenated (95% O_2_ and 5% CO_2_) modified artificial cerebrospinal fluid (ACSF) containing (in mM): KCl 2.5, NaH_2_PO_4_.H_2_O 1.2, NaHCO_3_ 30, D-glucose 25, HEPES 20, N-Methyl-D-glucamine 93, Na Ascorbate 5, Thiourea 2, Sodium pyruvate 3.0, N-acetyl-L-cystein 5.0, CaCl_2_.2H_2_O 0.5, and MgSO_4_.7H_2_O 10 (pH 7.4 and 310–320 mOsm). Sagittal lumbar spinal cord slices (400 μm thick) from both the ipsilateral (to the capsaicin application) and contralateral sides, with at least one attached dorsal root (L3 or L4, 8–12 mm), were prepared using a vibratome (VT1200S, Leica, Germany). These slices were incubated for ∼1 h at 35°C in oxygenated ACSF that contained (in mM): NaCl 124, KCl 2.5, NaH_2_PO_4_.H_2_O 1.2, NaHCO_3_ 15, D-glucose 12.5, HEPES 5, CaCl_2_.2H_2_O 2, and MgSO_4_.7H_2_O 2 (pH 7.4 and 310–320 mOsm). The slices were transferred into an imaging chamber and perfused with oxygenated ACSF at a rate of 10 ml/min throughout Ca^2+^ imaging at room temperature (22–25°C).

#### Ex vivo Ca^2+^ imaging

The offspring of SST^Cre^ x Ai95D and GAD2^Cre^ x Ai95D crossbreeding were used for Ca^2+^ imaging of SST-positive excitatory and GAD2-positive inhibitory interneurons in the superficial dorsal horn, as described previously^32^. The GCaMP6f green fluorescent signal from these neurons was detected using a microscope (BX51W1, Olympus, Japan) equipped with an ORCA Flash4.0LT camera (Hamamatsu, Japan) and PRIOR Lumen 300 LED light source (Cambridge, UK). A series of 512 x 512-pixel fluorescence images was acquired with HCImage software (Hamamatsu) at a 2-Hz sampling rate using a 40X objective immersed in ACSF. A suction electrode was used for electrical stimulation of the dorsal root at increasing intensities (10, 20, 50, 100, 300, and 500 μA; 0.5 ms pulse) to incrementally activate low- to high-threshold afferents. The dorsal root stimulation-evoked Ca^2+^ transients from the ipsilateral and contralateral spinal cords were recorded.

ImageJ (version 1.53v, NIH) was used to analyze all image files. Individual neurons showing changes in fluorescence upon dorsal root stimulation were identified, and the degree of change (ΔF) from the baseline fluorescence (F_0_) was divided by F_0_ (ΔF/F_0_). When comparing dorsal root stimulation-evoked Ca^2+^ transients in the ipsilateral dorsal horn neurons among different groups, their ΔF/F_0_ values were normalized to the averaged maximum ΔF/F_0_ in the corresponding contralateral slice.

### RNAscope

For RNAscope fluorescence in situ hybridization, dorsal root ganglia (DRG) were harvested from GPR37^fl/fl^ and TRPV1^Cre^GPR37^fl/fl^ mice. A modified protocol was used, adapted from the Advanced Cell Diagnostics (ACD) RNAscope Multiplex Fluorescent Reagent Kit v2 user manual (UM 323100, 2022). Predesigned probes for mouse *Gpr37* (Cat. # 319291) and *Trpv1* (Cat. # 313331-C3) were used.

### qRT-PCR

DRGs were harvested from GPR37^fl/fl^ and TRPV1^cre^GPR37^fl/fl^ mice and flash frozen on dry ice. Samples were stored at –80C° until RNA extraction. Tissues were homogenized using 0.5 mm glass beads and RNA extracted. RNA concentration was measured using a Nanodrop 2000C (ThermoFisher Scientific) according to the manufacturer’s instructions. cDNA was synthesized using QuantiTect Reverse transcription kit (QIAGEN, Cat no. 205311). qRT-PCR was done using SYBR green universal master mix (ThermoFisher Scientific, Cat no. 4309155), according to the manufacturer’s instruction with three technical replicates per sample (average of technical replicates reported) using primer pairs (Integrated DNA Technologies):

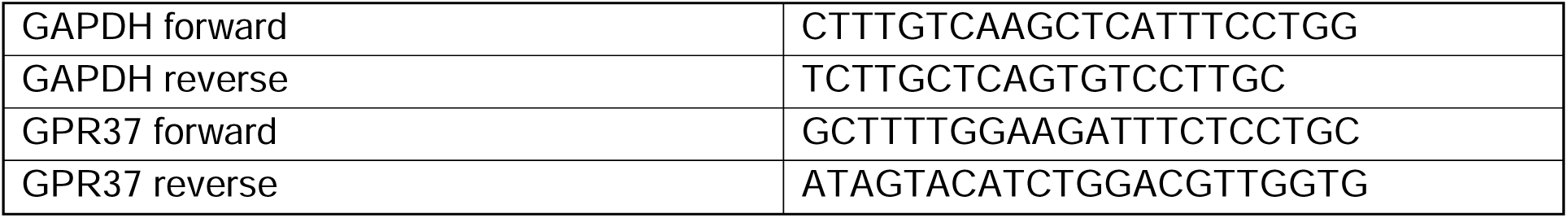

As a readout of gene transcript amount, we first calculated a difference in quantification cycle number (ΔCq) between GAPDH and GPR37 gene transcripts. Next, we calculated ΔΔCq by subtracting each sample’s ΔCq from the mean ΔCq of GPR37^fl/fl^ DRG samples; the smaller the ΔΔCq, the lower the GPR37 gene expression in DRG.

### Statistical Analyses

Sample size was estimated using the parameters from preliminary studies for 80% statistical power at an alpha level of 0.05 using G*Power 3.1 (Germany). Data from behavioral experiments were presented as mean±standard error of the mean (SEM), with N, the number of mice. Percent-withdrawal data were analyzed using a generalized linear mixed model (GLMM) - a binary logistic regression using a binomial distribution with a logit link function. Compound symmetry covariance structure was used for repeated measures over time, and degrees of freedom were approximated using the Satterthwaite method. Post hoc pairwise comparisons (vs the vehicle control) of estimated marginal means on the response scale (i.e., probability) were made using sequential Sidak tests. Withdrawal-latency data were analyzed using 2-way repeated measures (RM) ANOVA followed by Dunnett’s tests for pairwise comparisons (vs the vehicle control). qRT-PCR data were analyzed using a two-tailed unpaired t-test with Welch’s correction. Normalized calcium transients are presented as mean±SEM, with n, the number of neurons, and N, that of mice. Data from multiple neurons in each dorsal horn of individual mice were averaged per side to obtain the subject-level data representing each mouse and analyzed using 2-way RM ANOVA followed by Tukey’s tests. CPP data were analyzed using 1-way ANOVA followed by Tukey’s tests. GraphPad Prism 8 (Boston, MA, USA) and IBM® SPSS v28 (Chicago, IL, USA) were used for statistical analysis.

## Results

### Intrathecally injected TX14A and PD1 have no significant effect on normal nociception

Whether i.th. injection of GPR37 agonists affects normal nocifensive responses in naive mice (C57BL6N) was first determined. Three doses of TX14A (10, 25, and 50 μg) and two doses of PD1 (50 and 100 ng) were tested. Compared to the vehicle control group, all tested doses of i.th. TX14A (Fig 1) or PD1 (Fig 2) showed no significant effect on nocifensive withdrawal responses to mechanical (9.8-mN) and heat stimulation, suggesting that spinal-level GPR37 activation does not alter normal mechanical and heat nociception. The results also suggested that any changes in nocifensive behaviors in pain models following i.th. TX14A or PD1 are unlikely to be due to impaired withdrawal motor function.

**Fig 1.**
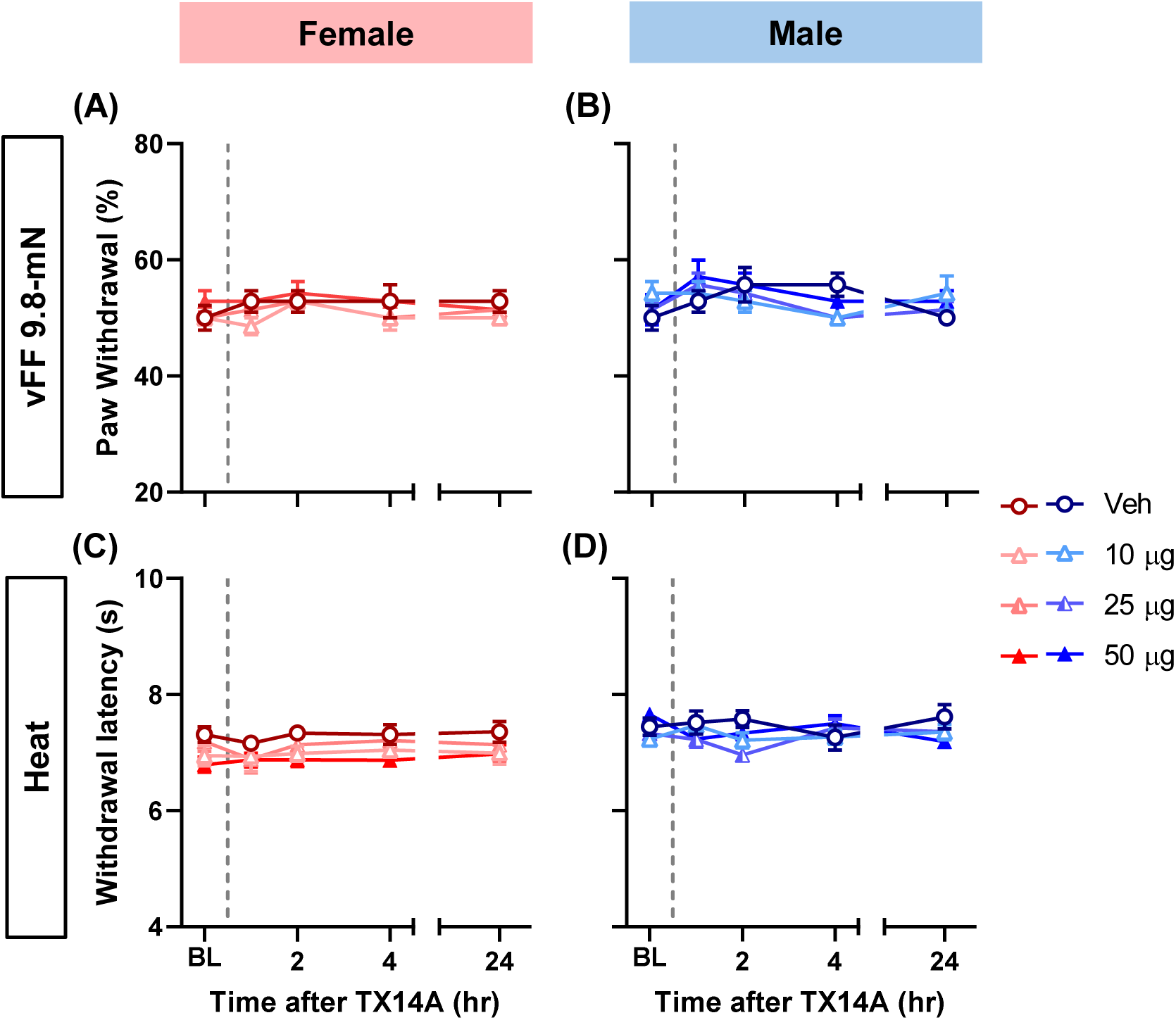
A single intrathecal (i.th.) injection of TX14A, a peptide agonist of GPR37, does not alter normal nociception. Three doses of TX14A (10, 25, and 50 μg; grey broken line) vs vehicle were tested (N=7/group). In both females (A, C) and males (B, D), there was no significant main effect (time or group) or interaction effect (time x group) on nocifensive withdrawal responses to 9.8-mN mechanical stimulation (A, B) or heat stimulation (C, D). BL, baseline.

**Fig 2.**
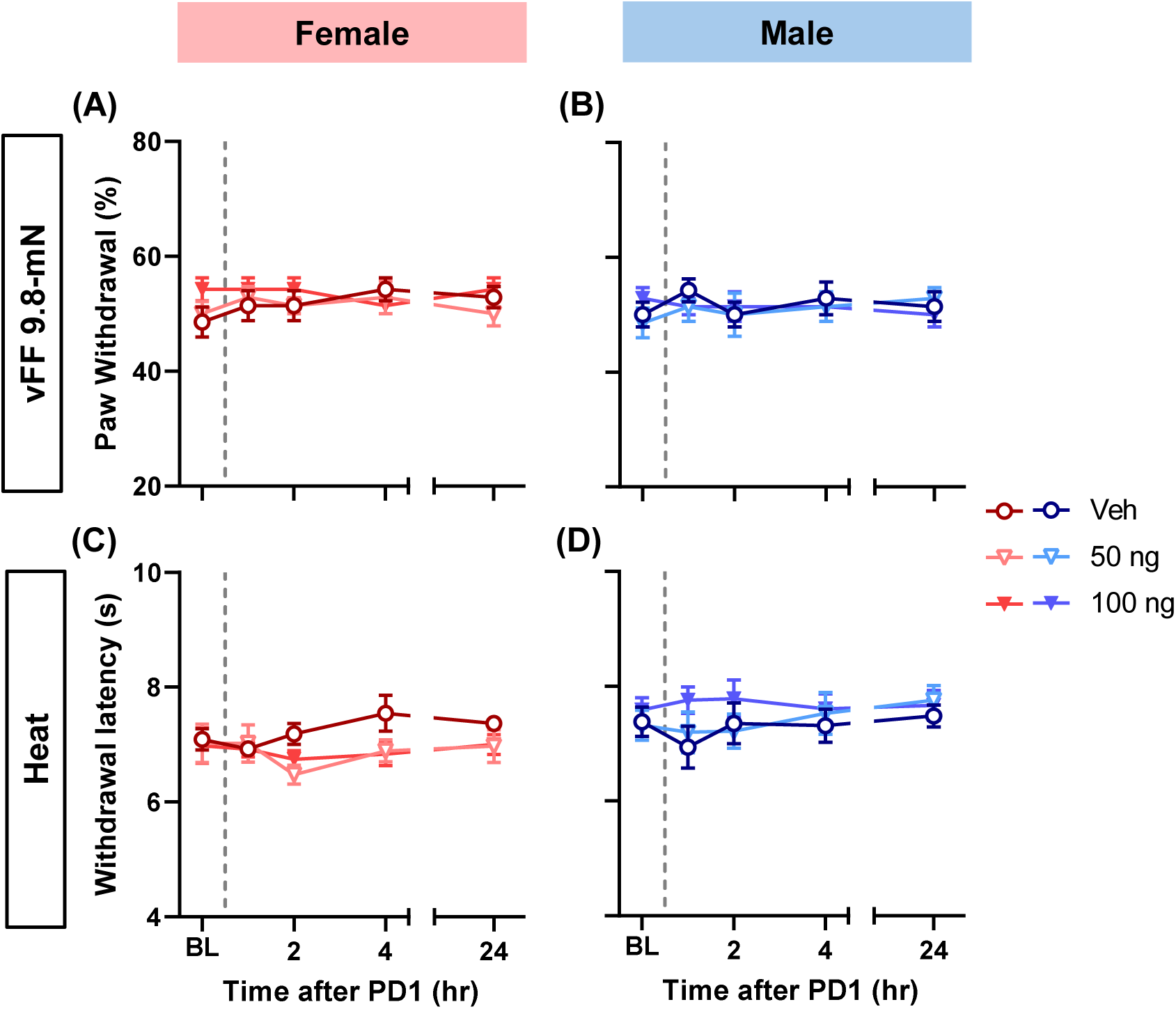
A single i.th. injection of protectin D1 (PD1), a lipid agonist of GPR37, does not alter normal nociception. Two doses of PD1 (50 and 100 ng; grey broken line) vs vehicle were tested (N=7/group). In both females (A, C) and males (B, D), there was no significant main effect (time or group) or interaction effect (time x group) on nocifensive withdrawal responses to 9.8-mN mechanical stimulation (A, B) or heat stimulation (C, D). BL, baseline.

### Spinal-level GPR37 activation inhibits capsaicin-induced mechanical and heat hypersensitivity

A single i.pl. capsaicin injection produces mechanical and heat hypersensitivity that persists for >24 hours in C57BL/6N mice^18,25^, with which long-term changes in the spinal nociceptive system are associated^19^. When intrathecally given 30 min after the capsaicin injection, TX14A dose-dependently inhibited the hypersensitivity in a biphasic manner; it produced a rapid-onset inhibition that gradually wore off, followed by a long-term effect still present at 24 h post-capsaicin in both sexes (Fig 3). TX14A (50 μg) was more effective against mechanical hypersensitivity in males than in females. Specifically, TX14A at this dose virtually resolved capsaicin-induced mechanical hypersensitivity in 24 hours in males but only partially attenuated it in females. However, no such sex difference was seen in the effects of TX14A on capsaicin-induced heat hypersensitivity.

**Fig 3.**
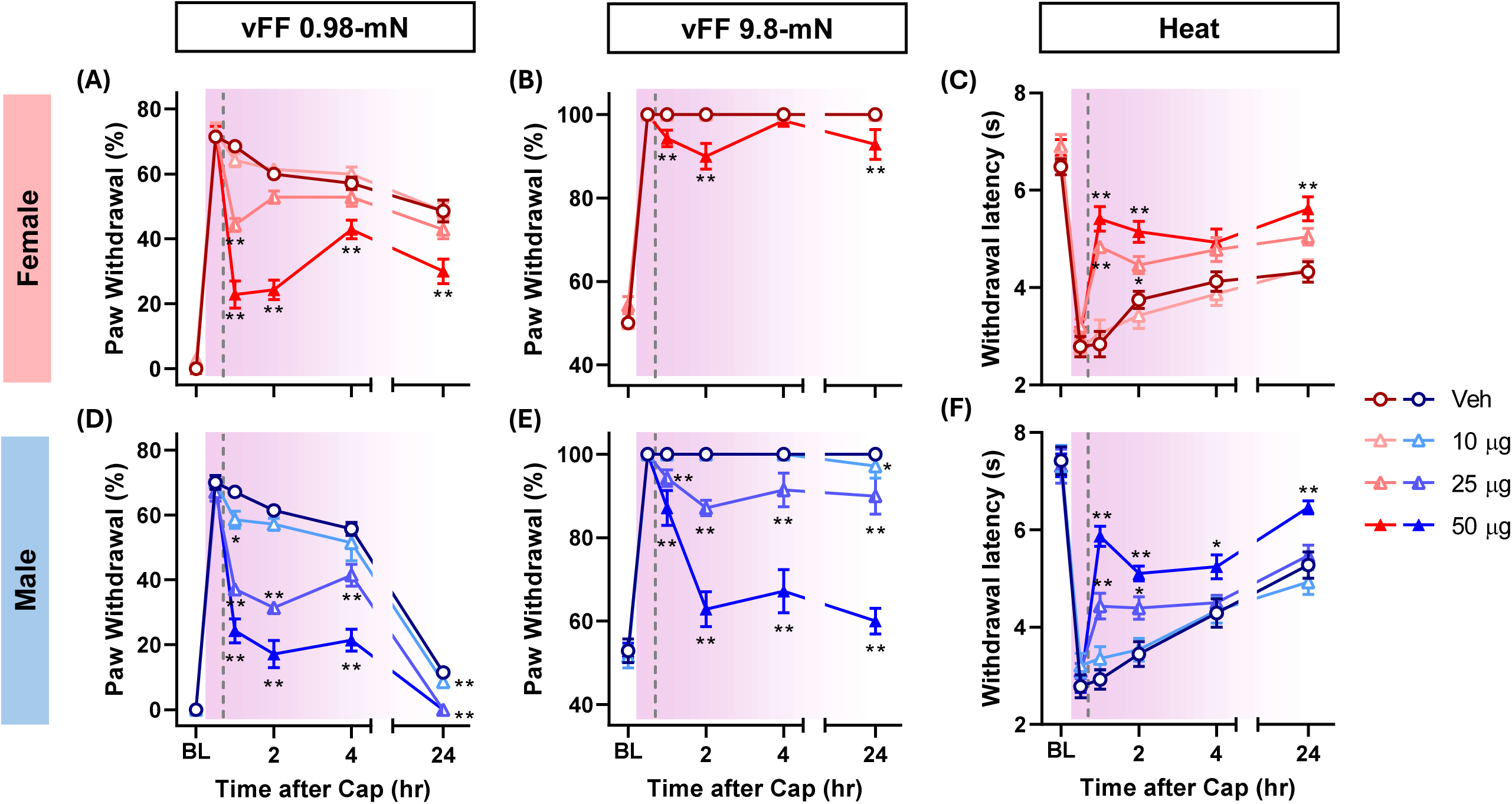
A single i.th. injection of TX14A dose-dependently inhibits capsaicin-induced mechanical and heat hypersensitivity both acutely and long-term. Intraplantar (i.pl.) injection of capsaicin produced mechanical (A, B, D, E) and heat (C, F) hypersensitivity that rapidly develops (in 30 min) and lasts for ≥24 hours in female (A-C) and male (D-F) C57BL/6N mice (purple-shaded area). The nocifensive withdrawal responses of these mice significantly differed among groups at each time point following drug administration (grey broken line), showing a dose-dependent biphasic inhibitory effect of i.th. TX14A on capsaicin-induced hypersensitivity. *p<0.05, **p<0.01 vs the vehicle group (Veh) by sequential Sidak tests following generalized linear mixed model (GLMM) analysis (A, B, D, E) or by Dunnett’s tests following 2-way RM ANOVA (C, F). N=7 per group. BL, baseline.

A single i.th. injection of PD1 also dose-dependently inhibited capsaicin-induced mechanical and heat hypersensitivity both acutely and long-term (Fig 4). As for the latter, 100 ng of PD1 virtually abolished mechanical hypersensitivity to 0.98-mN stimulation and partially inhibited that to 9.8-mN stimulation in both sexes. Overall, these findings indicated that spinal-level GPR37 activation resolves or persistently inhibits long-lasting mechanical and heat hypersensitivity in the capsaicin model.

**Fig 4.**
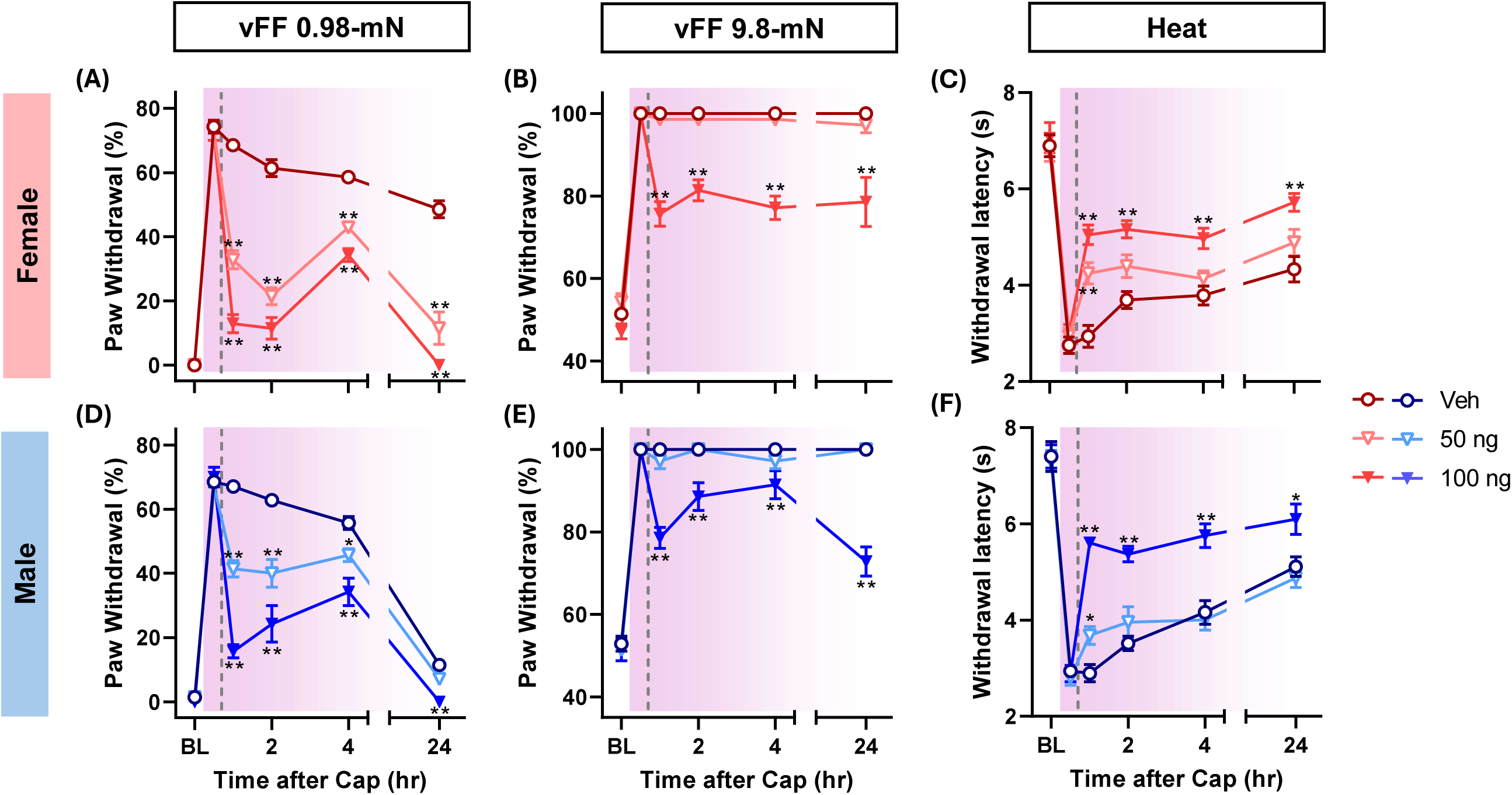
A single intrathecal (i.th.) injection of PD1 dose-dependently inhibits capsaicin-induced mechanical and heat hypersensitivity both acutely and long-term. I.pl. injection of capsaicin produced mechanical (A, B, D, E) and heat (C, F) hypersensitivity that rapidly develops (in 30 min) and lasts for ≥24 hours in female (A-C) and male (D-F) C57BL/6N mice (purple-shaded area). The nocifensive withdrawal responses of these mice significantly differed across groups at each time point following the drug administration (grey broken line), showing a dose-dependent biphasic inhibitory effect of i.th. PD1 on capsaicin-induced hypersensitivity. *p<0.05, **p<0.01 vs the vehicle group (Veh) by sequential Sidak tests following GLMM analysis (A, B, D, E) or by Dunnett’s tests following 2-way RM ANOVA (C, F). N=7 per group. BL, baseline.

### Spinal-level GPR37 activation prevents exaggerated nociception in the hyperalgesic priming model

Whether a single i.th. injection of TX14A affects mechanical and heat hypersensitivity development following i.pl. PGE_2_ injection in naïve (i.e., not-primed) mice was investigated. When given to naïve females two days prior to PGE_2_, TX14A (25 and 50 μg) slightly but significantly reduced PGE_2_-induced mechanical, but not heat, hypersensitivity at 4 h post-PGE_2_, whereas it did not affect PGE_2_-induced hypersensitivity in naïve males. Next, the effects of i.th. TX14A in mice that had already been primed by i.pl. IL-6 injection was examined. Compared to the vehicle-treated primed mice that developed exaggerated mechanical and heat hypersensitivity following i.pl. PGE_2_ injection, TX14A-treated primed mice developed significantly attenuated hypersensitivity following PGE_2_ (Fig 5).

**Fig 5.**
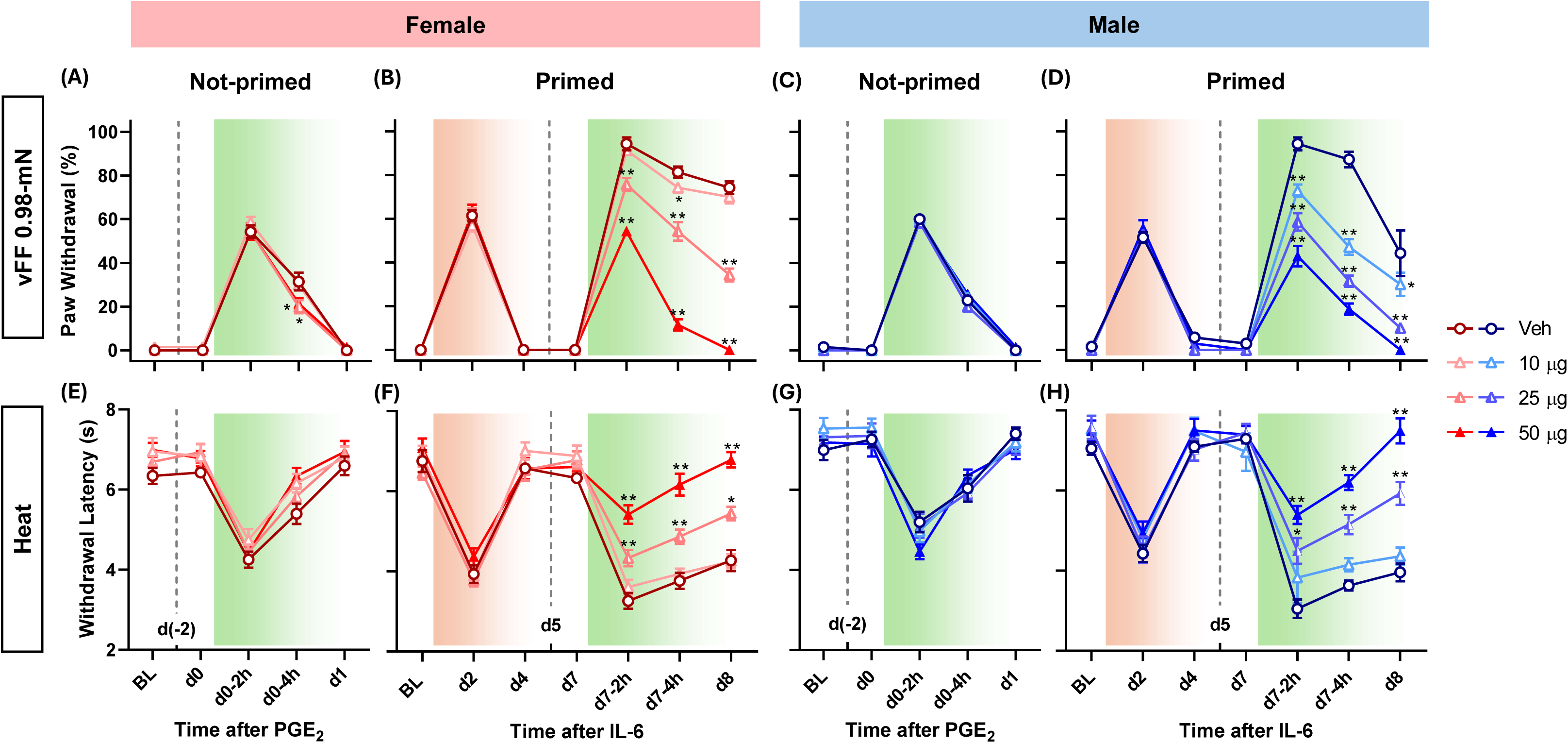
A single i.th. injection of TX14A dose-dependently erases hyperalgesic priming. When intrathecally given to naïve, not-primed females 2 days before i.pl. PGE_2_ injection (A, E), TX14A at 25 μg and 50 μg (grey broken line) reduced the degree of mechanical, but not heat, hypersensitivity at 4 h post-PGE_2_ (green-shaded area), unlike in naïve males (C, G). Mice that had been primed by i.pl. injection of interleukin-6 (IL-6, orange-shaded area) developed exaggerated mechanical (B, D) and heat (F, H) hypersensitivity in response to a subsequent i.pl. PGE_2_ injection. When TX14A was intrathecally given to primed mice 2 days before the i.pl. PGE_2_ injection, TX14A dose-dependently prevented the exaggeration of PGE_2_-induced hypersensitivity in both sexes (B, D, F, H). *p<0.05, **p<0.01 vs the vehicle group (Veh) by sequential Sidak tests following GLMM (A-D) or by Dunnett’s tests following 2-way RM ANOVA (E-H). N=7 per group. BL, baseline.

When intrathecally given to naïve females 2 days prior to PGE_2_, 100 ng of PD1 significantly decreased PGE_2_-induced heat, but not mechanical, hypersensitivity, whereas it did not alter PGE_2_-induced hypersensitivity in naïve males. However, further analysis revealed that this female-specific decrease likely resulted from the difference at baseline (BL) in this cohort of mice; specifically, when individual BL values were subtracted from the dataset, no significant difference was detected between groups in naïve females (Suppl Fig 1).

In primed mice, regardless of sex, both 50 and 100 ng of PD1 dose-dependently prevented the exaggeration of PGE_2_-induced mechanical and heat hypersensitivity (Fig 6). These results collectively suggested that spinal-level GPR37 activation erases the nociceptive system sensitization underlying hyperalgesic priming.

**Fig 6.**
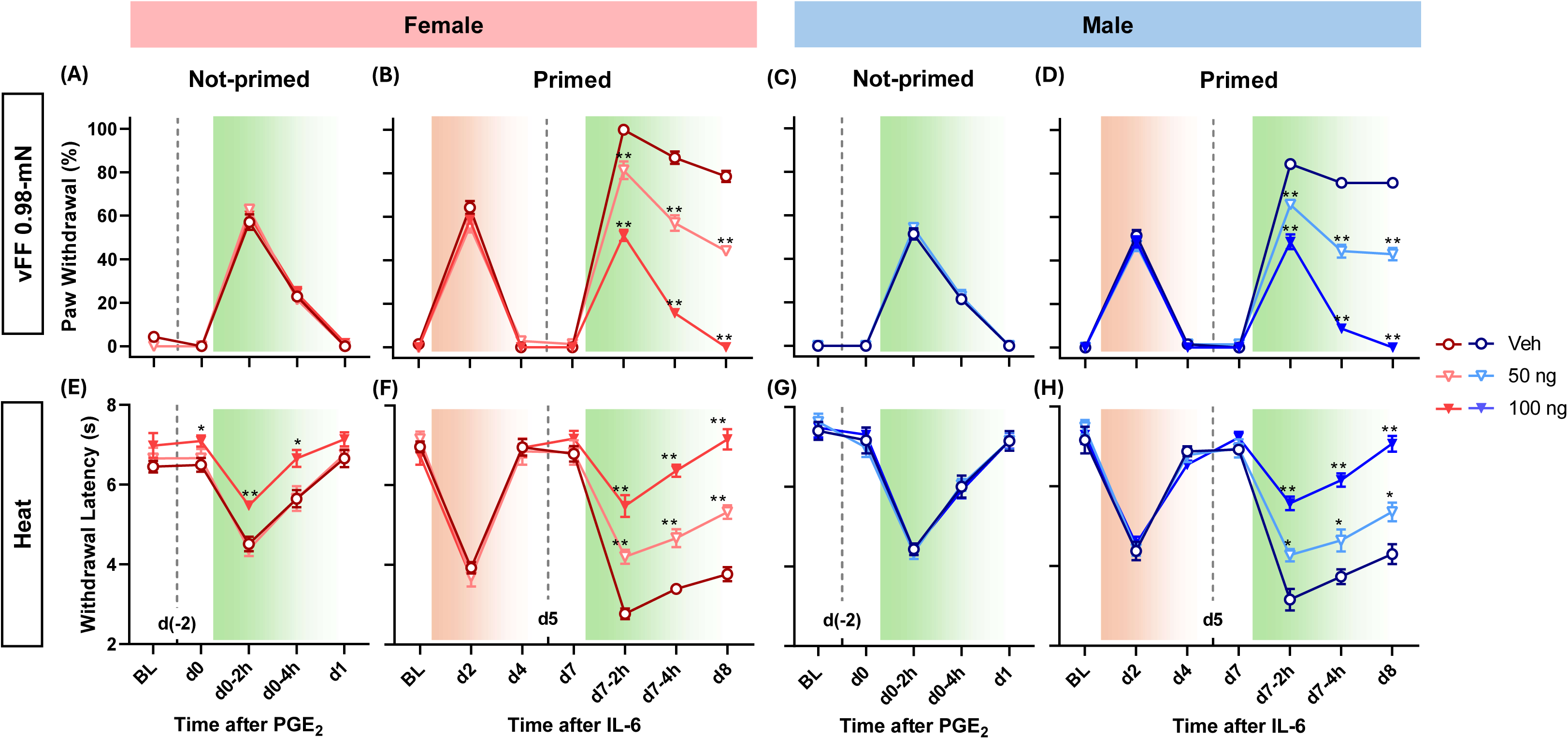
A single i.th. injection of PD1 dose-dependently erases hyperalgesic priming. When intrathecally given to not-primed females 2 days before i.pl. PGE_2_ injection (A, E), PD1 at 100 ng (grey broken line) reduced the degree of PGE_2_-induced heat, but not mechanical, hypersensitivity (green-shaded area), unlike in the male counterparts (C, G). Mice that had been primed by i.pl. injection of IL-6 (orange-shaded area) developed exaggerated mechanical (B, D) and heat (F, H) hypersensitivity in response to a subsequent i.pl. PGE_2_ injection. When PD1 was intrathecally given to primed mice 2 days before the i.pl. PGE_2_ injection, PD1 dose-dependently prevented the exaggeration of PGE_2_-induced hypersensitivity in both sexes (B, D, F, H). *p<0.05, **p<0.01 vs the vehicle group (Veh) by sequential Sidak tests following GLMM analysis (A-D) or by Dunnett’s tests following 2-way RM ANOVA (E-H). N=7 per group. BL, baseline.

### Global knockout of GPR37 prevents long-term inhibition of capsaicin-induced mechanical hypersensitivity and unpriming by i.th. TX14A

While PD1 was shown to be unable to activate GPR37-like 1 (GPR37L1), a close relative to GPR37^3^, TX14A nonselectively activates both GPR37 and GPR37L1^27,29^. Therefore, to determine whether the above-observed long-term inhibitory and unpriming effects of i.th. TX14A were mediated specifically by GPR37, earlier experiments were repeated in GPR37 global knockout (KO) mice and their wild-type (WT) genotypic control, focusing on mechanical hypersensitivity. Western blot of GPR37 in the spinal cord samples confirmed the absence of the receptor in GPR37 global KO mice (Suppl Fig 2).

In the capsaicin model (Fig 7), the effect of i.th. TX14A (50 μg) in the WT control was comparable to that seen in C57BL/6N mice, also showing a sex difference in its long-term efficacy against mechanical hypersensitivity. In GPR37 global KO mice, by contrast, TX14A failed to inhibit capsaicin-induced mechanical hypersensitivity long-term; it only produced a short-lasting, acute inhibition of mechanical hypersensitivity to 0.98-mN stimulation.

**Fig 7.**
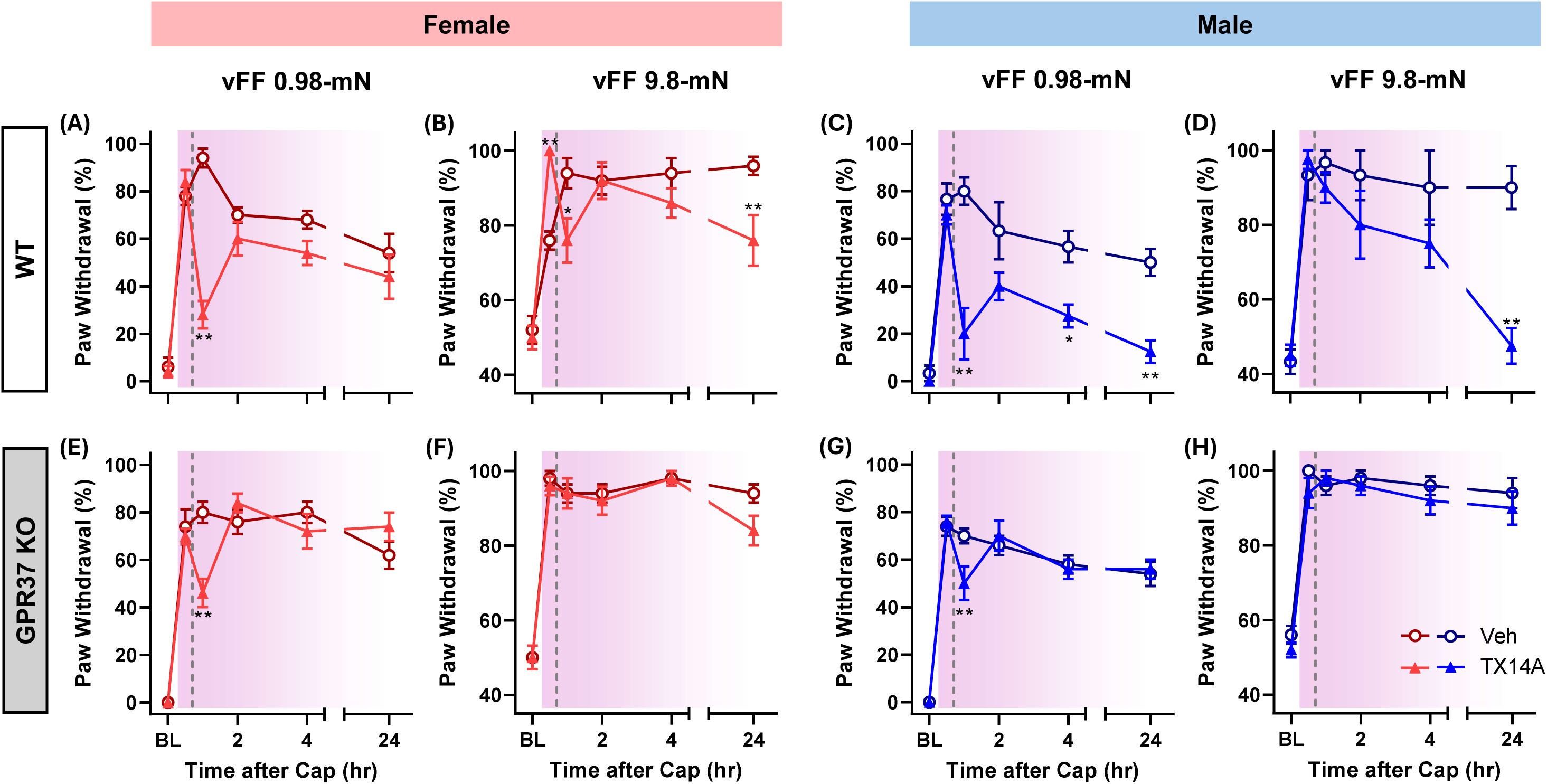
In the absence of GPR37, i.th. injection of TX14A fails to inhibit capsaicin-induced mechanical hypersensitivity long-term. In the wild-type (WT) control (A-D; N=3-5/group), TX14A (50 μg; grey broken line) inhibited capsaicin-induced mechanical hypersensitivity (purple-shaded area) both acutely and long-term. However, in the GPR37 knockout (KO) mice (E-H; N=5/group), TX14A only acutely inhibited hypersensitivity to 0.98-mN stimulation without producing a long-term (24 h post-capsaicin) effect. *p<0.05, **p<0.01 vs the vehicle group (Veh) by sequential Sidak tests following GLMM analysis. BL, baseline.

Compared to C57BL/6N mice (Figs 5 & 6), the WT control developed augmented mechanical hypersensitivity following i.pl. PGE_2_ injection even in a not-primed condition. However, the impact of hyperalgesic priming was still manifested as prolongation of PGE_2_-induced mechanical hypersensitivity in these mice (Fig 8). When intrathecally given after i.pl. IL-6 had primed the nociceptive system (i.e., 5 days after IL-6; 2 days before PGE_2_), TX14A effectively prevented such prolongation in the WT control but not in GPR37 global KO mice. Collectively, these findings indicated that GPR37 mediates the long-term inhibitory and unpriming effects of i.th. TX14A on enhanced nociception associated with nociceptive system sensitization.

**Fig 8.**
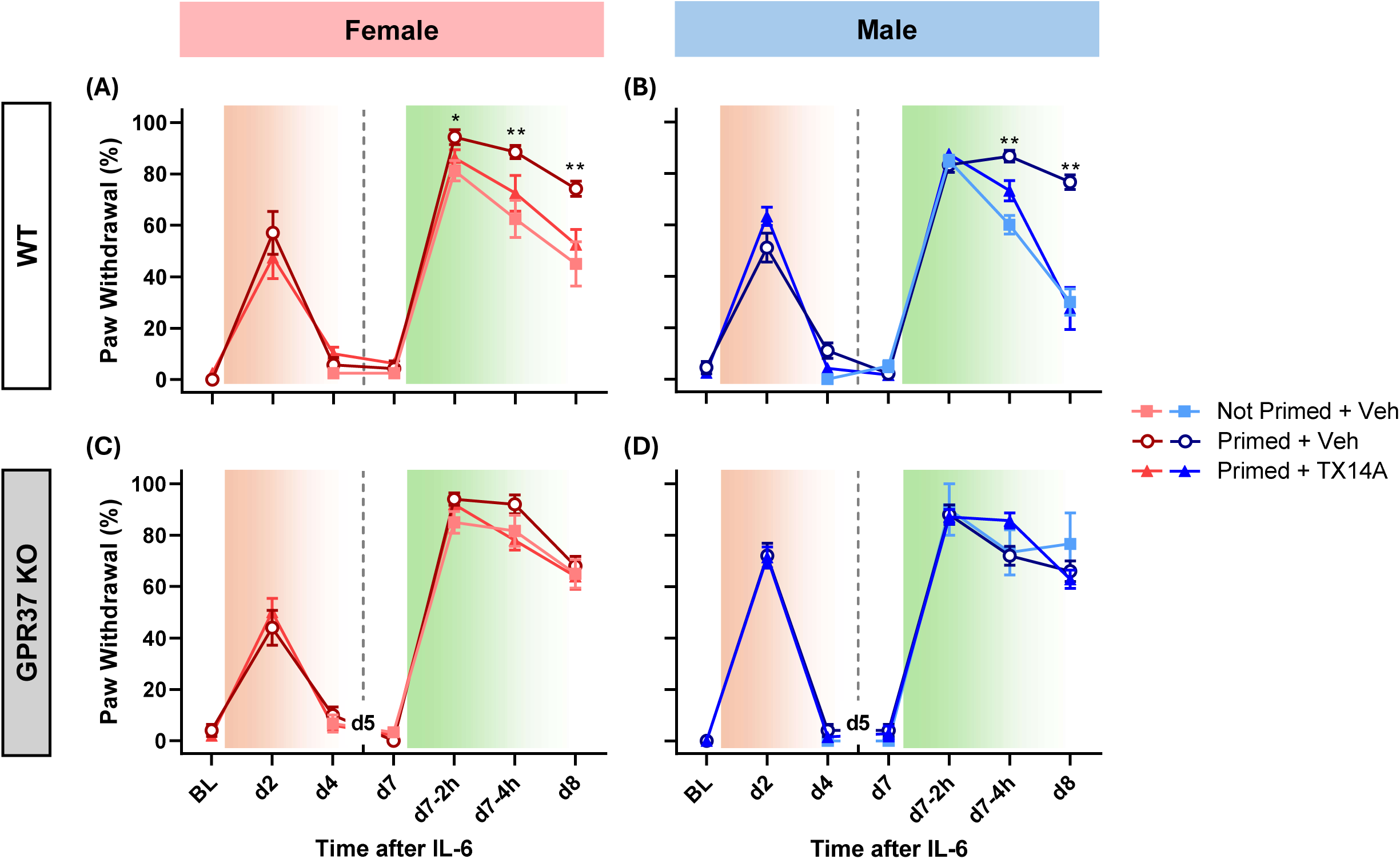
In the absence of GPR37, i.th. injection of TX14A fails to erase hyperalgesic priming. In the WT genotypic control (A, B; N=7-8/group), IL-6 (orange-shaded area)-produced hyperalgesic priming primarily manifested as prolonged mechanical hypersensitivity following i.pl. PGE_2_ (green-shaded area). When intrathecally given to primed WT control mice 2 days before i.pl. PGE2 injection, TX14A (50 μg; grey broken line) prevented such prolongation. However, in the GPR37 knockout (KO) mice (C, D; N=3-7/group), i.th. TX14A had no effect. *p<0.05, **p<0.01 vs the not-primed group by sequential Sidak tests following GLMM analysis. BL. Baseline.

### Conditional knockout of GPR37 in TRPV1-lineage DRG neurons prevents resolution of capsaicin-induced mechanical hypersensitivity and unpriming by i.th. TX14A

We next determined which specific cell type expressing GPR37 mediates the effects. GPR37 is highly expressed in presumed nociceptor populations in DRG^40^, suggesting that at the spinal level, their central terminals within the dorsal horn or cell bodies within DRG may be the action target of intrathecally administered GPR37 agonists. As these DRG neurons also highly express Transient receptor potential vanilloid 1 (TRPV1)^40^, we sought to determine whether TRPV1-positive DRG neurons expressing GPR37 are the specific cellular population mediating the spinal-level GPR37 agonism-triggered erasure of nociceptive system sensitization.

To this end, we selectively knocked out GPR37 in TRPV1-lineage neurons using Cre-lox gene recombination, producing TRPV1^Cre^GPR37^fl/fl^ (GPR37 cKO) mice. In the DRG of these mice, *Gpr37* mRNAs in *Trpv1* mRNA-positive DRG neurons were sparse, compared to GPR37^fl/fl^ mice (GPR37 flox, Fig 9A). qRT-PCR showed significantly reduced levels of GPR37 gene expression in DRG harvested from GPR37 cKO mice, compared to their genotypic control (Fig 9B). Together, these data confirmed the effective knockout of GPR37 in TRPV1-lineage DRG neurons.

**Fig 9.**
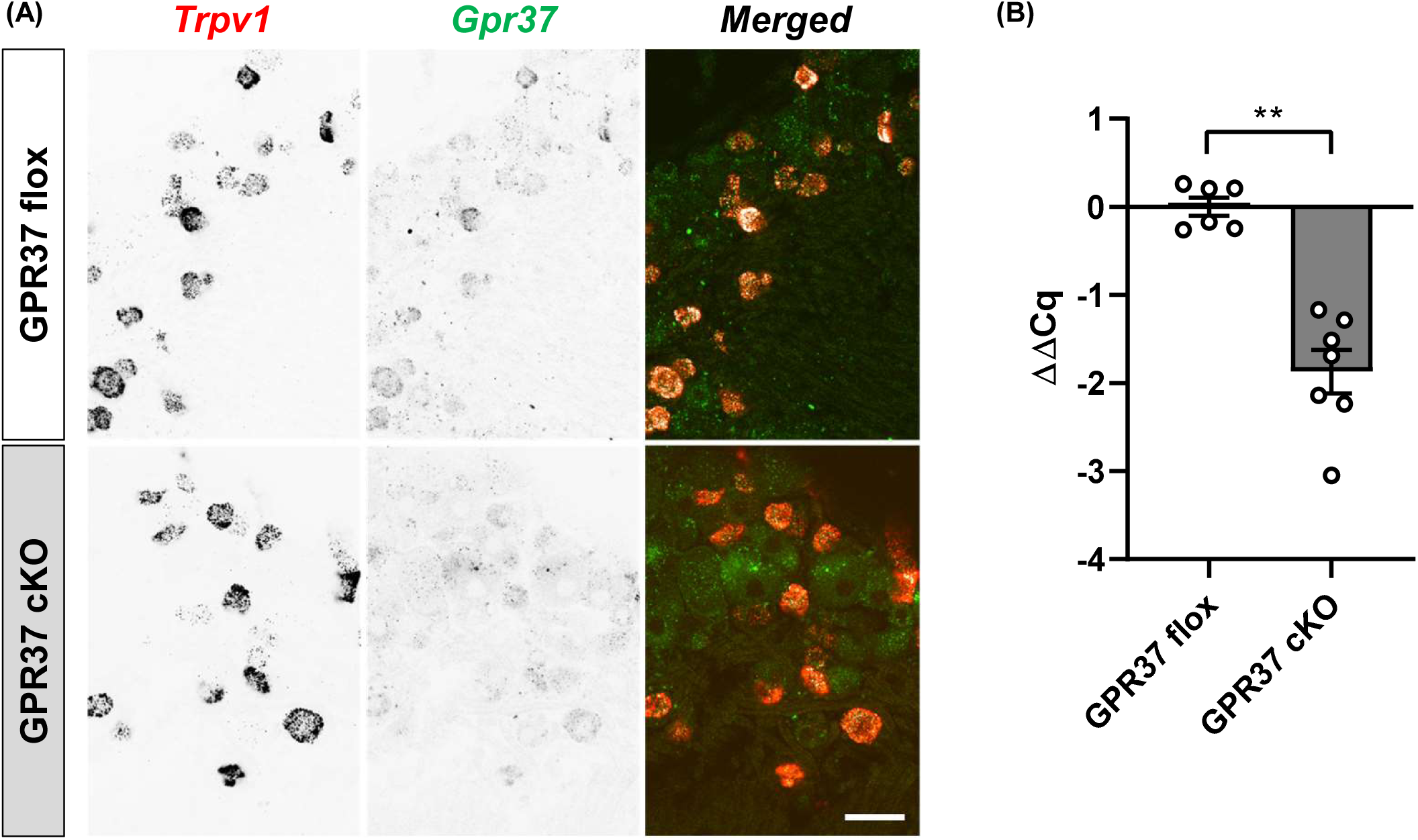
Validation of GPR37 conditional knockout in TRPV1-positive DRG neurons. (A) Representative RNAscope images showing the sparse expression of G*pr37* mRNAs in TRPV1-expressing DRG neurons in GPR37 conditional knockout (GPR37 cKO; i.e., TRPV1^Cre^GPR37^fl/fl^) mice, compared to the genotypic control, GPR37 flox (GPR37^fl/fl^) mice. Images of each target gene transcript were inverted to gray scales for visual clarity. Pixels containing dual signals (red: *Trpv1*, green: *Gpr37*) in merged images were colored in white. Bar=40 μm. (B) qRT-PCR showed significantly reduced gene expression of GPR37 in DRG of GPR37 cKO mice, compared to that in GPR37 flox mice. Data are presented as mean±SEM. N=6-7. **p<0.01 by two-tailed unpaired t-test with Welch’s correction.

In the capsaicin model, GPR37 flox mice (Fig 10A-D) developed allodynia- and hyperalgesia-like mechanical hypersensitivity that spontaneously resolved in 2 weeks. A single i.th. injection of TX14A produced a biphasic effect on the hypersensitivity: an acute, transient inhibition followed by an accelerated resolution. Notably, there was no apparent sex difference in the efficacy of i.th. TX14A (50 μg) to resolve long-term mechanical hypersensitivity in these mice. In GPR37 cKO mice, capsaicin-induced mechanical hypersensitivity persisted, and i.th. TX14A failed to inhibit or resolve it (Fig 10E-H). These data suggested that GPR37 in TRPV1-lineage DRG neurons is critical not only for the gradual, spontaneous erasure of capsaicin-induced sensitization of the nociceptive system but also for its facilitated erasure by spinal-level GPR37 activation.

**Fig 10.**
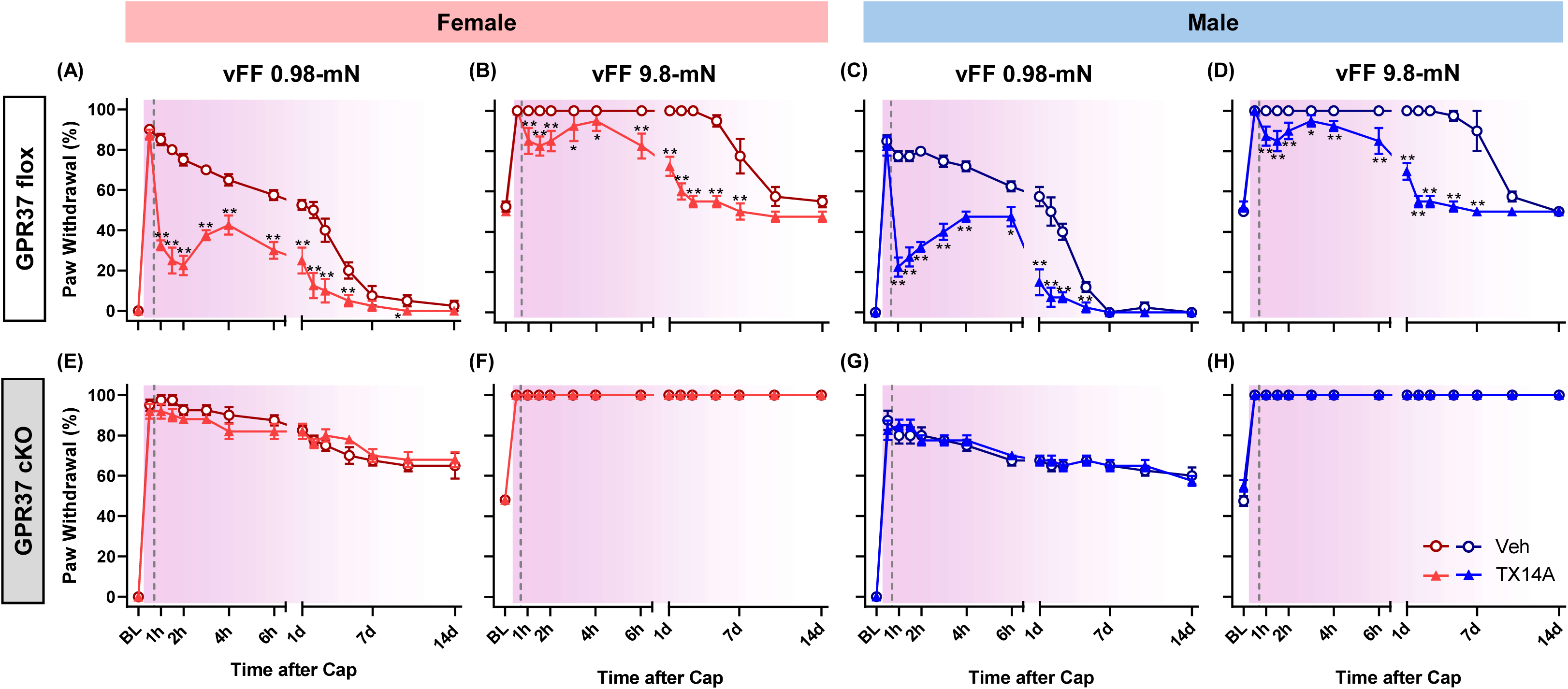
Capsaicin-induced mechanical hypersensitivity persists and does not resolve upon i.th. injection of TX14A in the absence of GPR37 in TRPV1-lineage DRG neurons. In GPR37 flox mice (A-D), a single i.th. injection of TX14A showed a biphasic effect on capsaicin-induced mechanical hypersensitivity in both sexes: an acute, transient inhibition followed by an accelerated resolution. In contrast, hypersensitivity persisted in GPR37 cKO mice, and i.th. TX14A failed to trigger resolution (E-H). N=4-5 in each group. Data are presented as mean±SEM. *p<0.05, **p<0.01 vs the vehicle (Veh) group by sequential Sidak tests following GLMM analysis. BL, baseline.

In the hyperalgesic priming model, the duration and intensity of mechanical hypersensitivity following i.pl. IL-6 injection was significantly increased in GPR37 cKO mice (Fig 11A & D). Primed GPR37 flox mice developed exaggerated mechanical hypersensitivity following i.pl. PGE_2_ injection, which was completely prevented by a single i.th. TX14A (50 μg) given after the resolution of IL-6-induced mechanical hypersensitivity (Fig 11B & E). However, in GPR37 cKO mice, i.th. TX14A failed to unprime the nociceptive system (Fig 11C & F). Collectively, these findings indicated that GPR37 in TRPV1-lineage DRG neurons plays a key role in the erasure of acute injury-induced sensitization of the nociceptive system.

**Fig 11.**
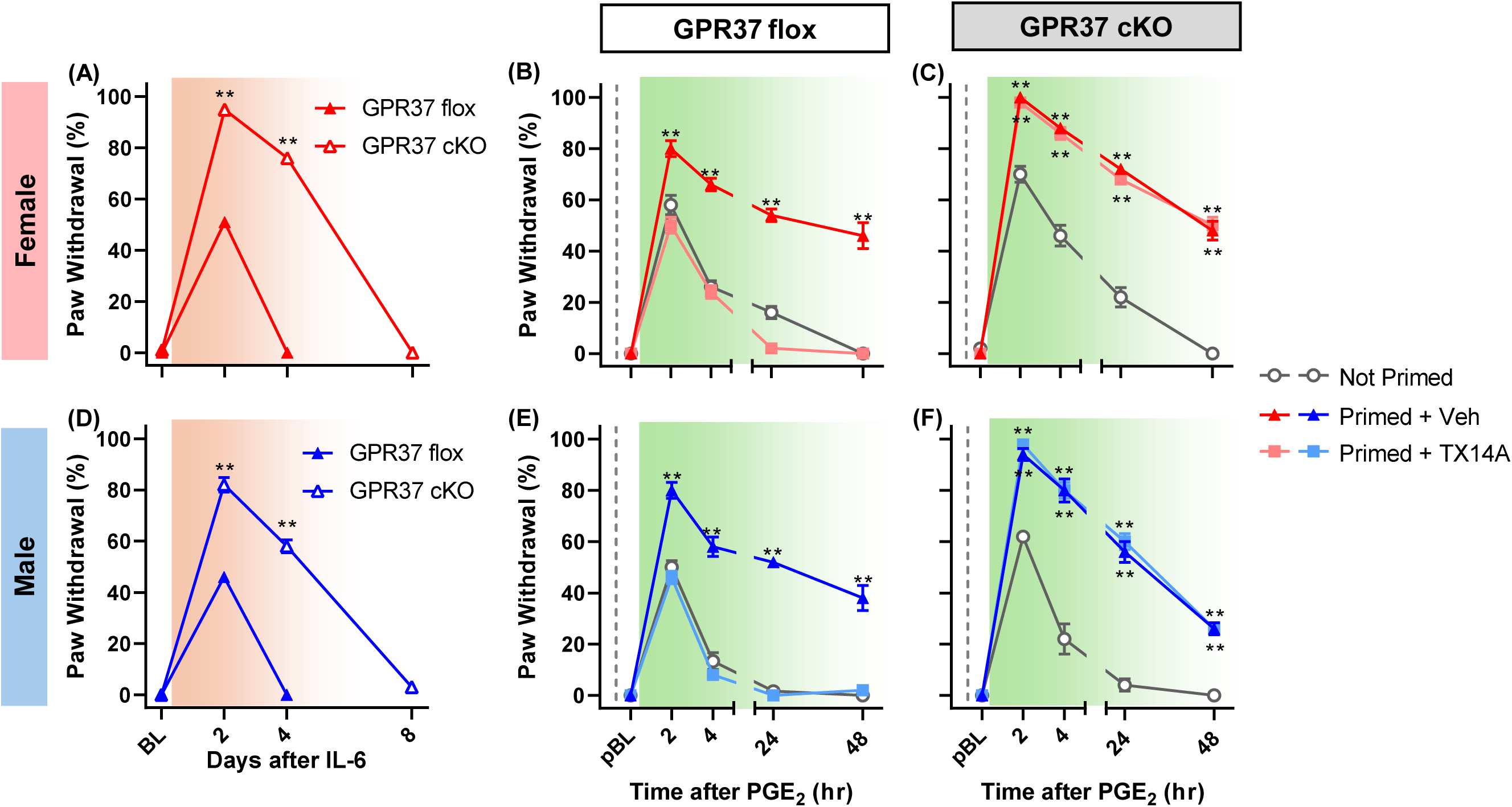
Intrathecally injected TX14A does not erase hyperalgesic priming in the absence of GPR37 in TRPV1-lineage DRG neurons. An i.pl. injection of IL-6 produced greater mechanical hypersensitivity in GPR37 cKO mice than in GPR37 flox mice (A & D). When intrathecally given in primed GPR37 flox mice, TX14A unprimed the nociceptive system, preventing the exaggeration of PGE_2_-produced mechanical hypersensitivity (B & E). In GPR37 cKO mice, however, i.th. TX14A failed to unprime the nociceptive system (C & F). N=4-5 in each group. Data are presented as mean±SEM. **p<0.01 vs GPR37 flox (in A & D) or the not-primed group (in B, C, E, and F) by sequential Sidak tests following GLMM analysis. BL, baseline. pBL, primed baseline.

### Spinal-level GPR37 activation rescues dorsal horn neurons from capsaicin-induced long-term changes

To support the behavioral evidence that spinal-level GPR37 activation erases nociceptive system sensitization, we next investigated whether i.th. TX14A abrogates capsaicin-induced long-term changes in neuronal responsiveness to afferent inputs in the superficial dorsal horn (SDH). To this end, transgenic mice expressing GCaMP6f either in SST-positive (SST^Cre^Ai95D) or GAD2-positive neurons (GAD2^Cre^Ai95D) were used. In the SDH, 82±2% of neurons labeled using the SST^Cre^ line express SST gene transcripts^10^, and 86±3% of neurons labeled using the GAD2^Cre^ line are immunoreactive for PAX2, a marker of inhibitory neurons in the SDH^32^. In these mice, to maximize the yield of sensitized dorsal horn neurons from each mouse in ex vivo slice preparations, the sciatic nerve was directly exposed to capsaicin (day 0). Afterward, TX14A (50 μg) or saline was intrathecally administered twice - immediately after the capsaicin application and on the following day. On day 7, mechanical hypersensitivity (to 9.8-mN stimulation) was assessed in both hind paws, and then the spinal cord was harvested (Fig 12A). The control group received the vehicle of capsaicin on the sciatic nerve and i.th. saline.

**Fig 12.**
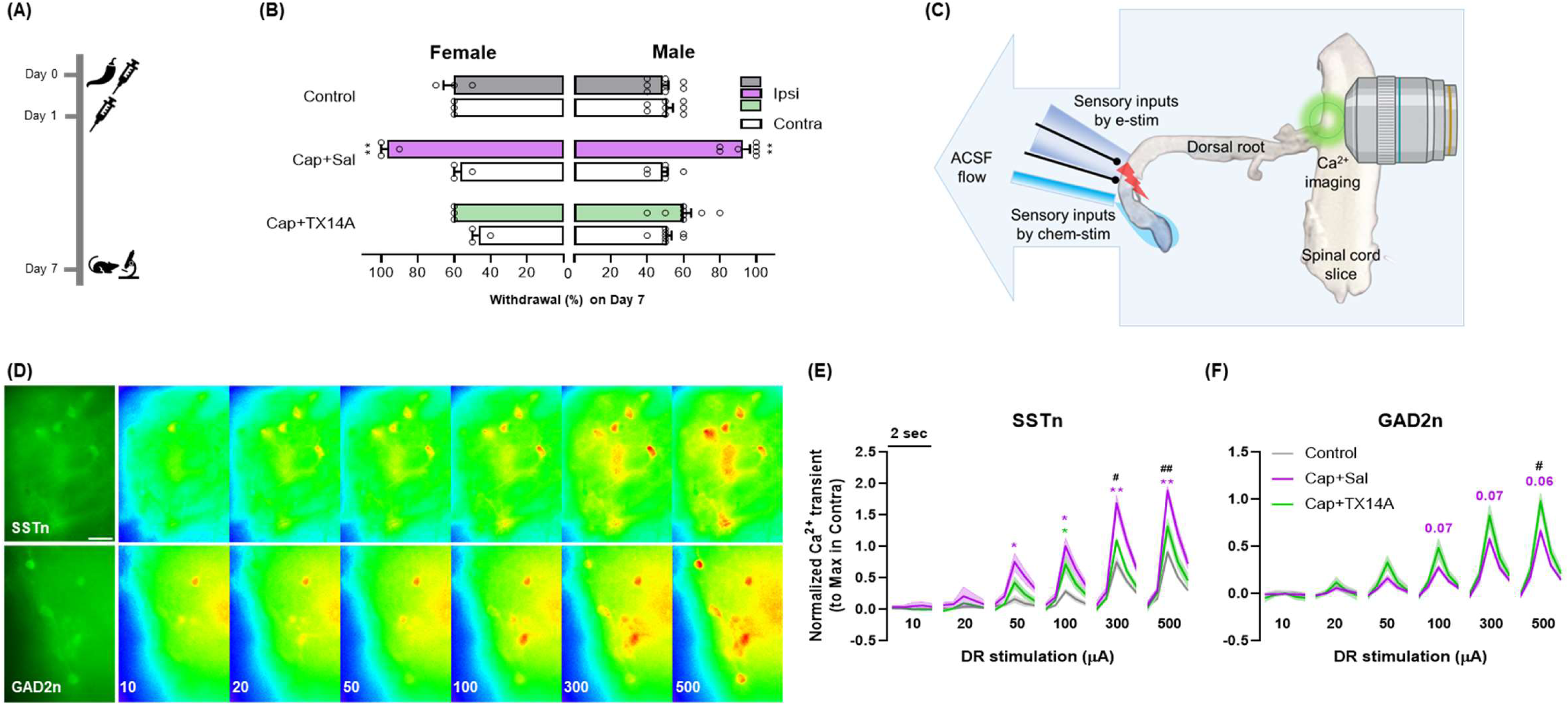
I.th. injection of TX14A resolves long-term changes in dorsal horn neuronal responsiveness to afferent inputs. (A) In mice that express GCaMP6f in somatostatin (SST)-positive excitatory interneurons (SSTn) or GAD2-positive inhibitory interneurons (GAD2n) in the dorsal horn, capsaicin (or its vehicle) was unilaterally applied to the sciatic nerve, and TX14A (vs saline) was intrathecally given twice (days 0 and 1). On day 7, following the measurement of nocifensive withdrawals from 9.8-mN mechanical stimulation, ex vivo Ca^2+^ imaging was performed. (B) Both in females (N=3/group) and males (N=6-8/group), the TX14A treatment (Cap+TX14A) resolved long-lasting mechanical hypersensitivity in the ipsilateral hind paw. **p<0.01 vs the contralateral paw by sequential Sidak tests following GLMM analysis. (C) Experimental set-up of ex vivo Ca^2+^ imaging. The attached dorsal root (DR) was electrically stimulated to generate afferent inputs to the dorsal horn. (D) Representative images of SSTn and GAD2n and their GCaMP6f signal strengths at each DR stimulation intensity. Bar=30 μm. (E) In SSTn, compared to DR stimulation-evoked Ca^2+^ transients in the ipsilateral dorsal horn of the control group (n=41, N=3), those were potentiated in the Cap+Sal group (n=71, N=4). The Cap+TX14A group (n=53, N=4) showed Ca^2+^ transients comparable to those in the control group. (F) In contrast, GAD2n in the ipsilateral dorsal horn of the Cap+Sal group (n=186, N=5) showed a trend toward decreased Ca^2+^ transients, compared to those in the control group (n=129, N=5), whereas the Cap+TX14A group (n=92, N=5) showed no difference from the control group; Ca^2+^ transients of both groups completely overlapped, indistinguishable in the figure. *p<0.05, **p<0.01 vs the control group; #p<0.05, ##p<0.01 between the Cap+Sal and Cap+TX14A groups by Tukey’s tests following 2-way RM ANOVA.

In the control group, there was no difference in mechanical sensitivity between the ipsilateral and contralateral hind paws on day 7. However, in mice treated with capsaicin and then i.th. saline (the Cap+Sal group), the ipsilateral hind paw remained hypersensitive to 9.8-mN stimulation compared to the contralateral hind paw. In contrast, in mice treated with capsaicin and then i.th. TX14A (the Cap+TX14A group), there was no significant mechanical hypersensitivity in the ipsilateral hind paw (Fig 12B). Notably, there was no sex difference in the efficacy of TX14A; two i.th. injections of TX14A effectively resolved hypersensitivity to 9.8-mN stimulation in both sexes. Therefore, in the subsequent Ca^2+^ imaging (Fig 12C-F), data obtained from females and males were pooled.

In the SDH harvested 7 days after the capsaicin application, dorsal root stimulation-evoked Ca^2+^ transients in SST-positive excitatory interneurons (SSTn) were different across groups (F(2,8)=8.977, p=0.009; Fig 12D & E). Post hoc analyses revealed that their responses to afferent inputs were overall potentiated in the Cap+Sal group, compared to those in the control group (t(8)=4.085, Tukey’s adjusted p=0.009). The Ca^2+^ transients of SSTn in the Cap+TX14A group were overall not different from those in the control group (t(8)=1.412, Tukey’s adjusted p=0.38) but significantly smaller than those in the Cap+Sal group (t(8)=2.888, Tukey’s adjusted p=0.048), suggesting that TX14A had rescued SSTn from capsaicin-induced sensitization. In line with this, when non-normalized Ca^2+^ transients were compared between ipsilateral and contralateral dorsal horns in each group, only the Cap+Sal group showed a trend (F(1,6)=4.943, p=0.068 by 2-way RM ANOVA) toward a difference in neuronal responsiveness to afferent inputs (Suppl Fig 3A-C).

As for Ca^2+^ transients in GAD2-positive inhibitory interneurons (GAD2n), there was a significant interaction of group x stimulation intensity (F(58,348)=2.349, p<0.0001; Fig 12D & F). Pairwise comparisons between groups at each stimulation intensity revealed that their responses to afferent inputs tended to be depressed (Tukey adjusted p=0.058-0.071) at high intensities (≥100 μA) in the Cap+Sal group, compared to those in the control group. Notably, at 500 μA stimulation, the Ca^2+^ transient of the Cap+TX14A group was significantly greater than that of the Cap+Sal group (t(5.081)=4.748, Tukey’s adjusted p=0.044), suggesting that TX14A had normalized capsaicin-induced long-term disinhibition. In line with this, when non-normalized Ca^2+^ transients were compared between ipsilateral and contralateral dorsal horns in each group, only the Cap+Sal group showed a significant difference (F(1,8)=7.448, p=0.026 by 2-way RM ANOVA) in neuronal responsiveness to afferent inputs (Suppl Fig 3D-F).

### Activation of GPR37 in the central nervous system has no apparent abuse liability

As the above findings collectively suggested spinally directed GPR37 agonists as novel pain therapeutics to erase pain hypersensitivity without affecting normal nociception, their safety in terms of abuse potential was evaluated. To this end, conditioned place preference (CPP) tests were performed using saline as the negative control, morphine (7-10 μg) as the positive control, and TX14A (50 μg) and PD1 (100 ng) as GPR37 agonists. As shown in Fig 13, mice conditioned with either GPR37 agonist did not develop place preference for the drug-paired chamber compared to the saline-treated negative control group, suggesting that centrally activating GPR37 agonists may have little to no abuse liability.

**Fig 13.**
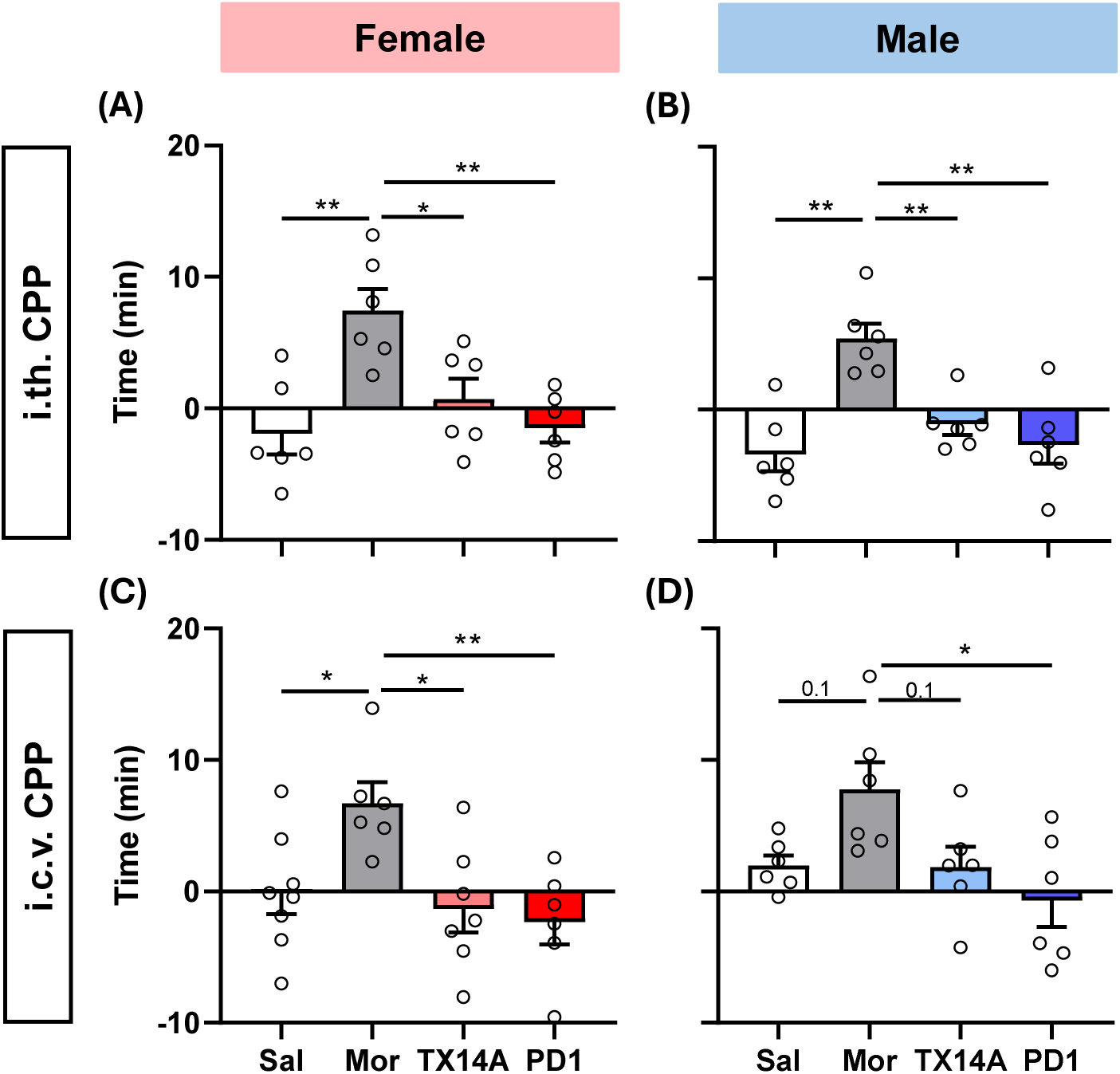
GPR37 agonism in the central nervous system does not show apparent abuse potential. Unlike morphine (Mor)-treated mice, TX14A- or PD1-treated mice did not develop conditioned place preference for the treatment-paired chamber, compared to the saline (Sal)-treated mice, after intrathecally (A, B; N=6/group) or intracerebroventricularly (C, D; N=6-8/group) receiving treatments for 3 consecutive conditioning days. *p<0.05, **p<0.01 by Tukey’s tests following 1-way ANOVA.

## Discussion

Intense nociceptive inputs induce long-term changes in the spinal nociceptive system, resulting in heightened and long-lasting pain. Aligning with prior findings^6,24,32^, the present results of ex vivo Ca^2+^ imaging demonstrate that such changes consist of neuron type-dependent long-term plasticity in opposing directions: the responsiveness to afferent inputs is potentiated in excitatory neurons but depressed in inhibitory neurons in the SDH. This study provides preclinical evidence that spinal-level GPR37 activation normalizes such long-term changes and thus holds potential as a novel therapeutic approach to resolving pain, instead of temporarily suppressing it. Importantly, GPR37 activation at the spinal level did not alter normal mechanical and heat nociception, suggesting that spinally delivered GPR37 agonists may selectively target maladaptive nociceptive processing in a pathological pain condition. This may be an ideal approach to treating pathological, persistent pain while preserving physiological pain’s protective role. In addition, centrally activating GPR37 did not show an apparent abuse liability in the CPP test, further supporting the translational potential of spinal-level GPR37 activation as a therapy for persistent pain.

Since TX14A is known to be rapidly degraded in the brain and not accumulated after repeated dosing^13,37,38^, the long-term effect of TX14A in the capsaicin model and hyperalgesic priming model seems unlikely to be due to the peptide persistently staying (>24 h) in the system. Instead, it suggests that TX14A may trigger a GPR37-mediated event that outlasts the lifespan of the peptide in the nervous system, which subsequently leads to the erasure of nociceptive system sensitization. The biphasic effects of i.th. TX14A in the capsaicin model aligns with this idea, and future pharmacokinetic studies will be necessary to determine whether the initial rapid-onset effect (which wears off in ∼3 hours) directly correlates with TX14A concentration within the spinal cord.

As for PD1, it was shown to be readily metabolized in an assay system using human hepatoma cells, with one of its metabolites retaining biological activity to inhibit neutrophil chemotaxis^1^. Therefore, although TX14A and PD1 commonly produced persistent inhibition or resolution of increased nociception in the capsaicin model (except for mechanical hypersensitivity in female C57BL6N mice, see below) and unpriming in the hyperalgesic priming model, differences in their metabolic fates and pharmacokinetic properties may account for any observed differences in their effects on nocifensive behaviors. Two potential putative binding sites on GPR37 have been identified using homology modeling: one for small molecules including PD1, and the other for TX14A^2^. Activating GPR37 through these distinct binding sites could also result in pharmacodynamic differences between PD1 and TX14A. In addition, unlike PD1, TX14A activates both GPR37 and GPR37L1, which may further contribute to differences in the effects of the two GPR37 agonists in pain models. Regarding this, it is noteworthy that TX14A still briefly inhibited capsaicin-induced hypersensitivity to 0.98-mN stimulation in GPR37 global KO mice. It would be interesting to examine in future studies if GPR37L1 mediates this small residual effect. GPR37L1 is primarily expressed in satellite glial cells in DRG^3,12^ and astrocytes in the spinal cord^44^, playing a critical role in alleviating neuropathic pain^3,44^.

Interestingly, our experiments using GPR37 cKO mice found no such residual acute effect of i.th. TX14A to attenuate allodynia-like mechanical hypersensitivity (Fig 13). This discrepancy could be explained by a potential compensatory mechanism (e.g., enhanced GPR37L1 signaling) due to “global” GPR37 KO, while such compensation may be absent in our “conditional” KO condition where GPR37 is still present in cells other than TRPV1-lineage ones.

A single i.th. injection of TX14A was less effective in inhibiting capsaicin-induced mechanical hypersensitivity in female than in male C57BL/6N mice. As female rodents and humans experience more intense and longer-lasting mechanical pain following capsaicin application^14,15,17,18^, a higher dose than 50 μg of TX14A may be necessary in females to produce effects comparable to those in males. This idea is supported by the qualitative observation that the effects of TX14A 50 μg on mechanical hypersensitivity in females (Fig 3A, B) resembled those of TX14A 25 μg in males (Fig 3D, E).

Speaking of the abovementioned sex difference in the efficacy of TX14A in the capsaicin model, it is noteworthy that no such difference was observed against capsaicin-induced ‘heat’ hypersensitivity in C57BL/6N mice (Fig 3C, F). Given that mechanical and heat nociception is mediated by two distinct afferent populations (MRGPRD-positive neurons for the former and TRPV1-positive ones for the latter) in the mouse^9^, these results suggest that capsaicin-induced long-term changes in processing the two distinct afferent inputs are mechanistically different, resulting in sensory modality-dependent efficacy of i.th. TX14A.

In addition, we observed an inconsistency in the sex-dependent efficacy of i.th. TX14A among mice of different genetic backgrounds in the capsaicin model. Unlike C57BL/6N mice and GPR37 global KO mice (congenic B6.129P2 strain), GPR37 flox mice, whose background strain is C57BL/6J, did not show a sex difference in the efficacy of a single i.th. injection of TX14A (50 μg) to inhibit/resolve capsaicin-induced mechanical hypersensitivity. There are established mouse strain^30^ and substrain differences in pain phenotypes. As for the latter, C57BL/6J mice show shorter withdrawal latency in hot plate and tail withdrawal assays^7,8^, while C57BL/6N mice show increased formalin-induced sensitivity to spontaneous pain-like behaviors and greater long-term mechanical hypersensitivity^39^. Therefore, we do not disregard the possibility that mouse strain and substrain differences in the nociceptive system responsiveness to injury may contribute to the sex difference in the efficacy of a single-dose TX14A in the capsaicin model.

With respect to the foregoing, we also did not observe such a sex difference in the experiments where TX14A (50 μg) was administered twice at a 24-h interval in SST/GAD2^Cre^Ai95D mice whose “genetic background consists of a mix of 3 or more inbred strains, contains outbred contributions, or contains unknown contributions,” according to the Jackson Laboratory. Therefore, the interpretation of no sex difference in the efficacy of i.th. TX14A in these modified capsaicin model experiments is constrained, as it could result from mouse strain differences or the additional dose of TX14A that could have negated potential sex-specific neural mechanisms in a single-dose condition.

We found that the loss of GPR37 in TRPV1-lineage DRG neurons impaired the resolution of both capsaicin-induced mechanical hypersensitivity and IL-6-induced hyperalgesic priming, indicating that GPR37 in this cellular population is the key to the erasure of nociceptive system sensitization. The downstream mechanisms of these findings require elucidation, particularly how GPR37 activation in these neuronal cell bodies (in DRG) and/or central terminals (within the dorsal horn) affects the neuronal activity in the DRG and dorsal horn both acutely and long-term. PD1 has been shown to instantly prevent capsaicin-evoked currents in DRG neurons and increased frequency of spontaneous excitatory postsynaptic currents in dorsal horn neurons^33^, suggesting that the acute inhibitory effect of spinal-level GPR37 agonism may result from inhibition of TRPV1-positive afferent activity. While the mechanisms underlying the long-term effect remain unknown, the restoration of the excitatory-inhibitory balance in the spinal nociceptive circuit seems to constitute the GPR37-triggered erasure of nociceptive system sensitization, as our Ca^2+^ imaging results demonstrate that i.th. TX14A has normalized capsaicin-induced long-term changes in afferent-evoked activation of both excitatory and inhibitory interneurons in the SDH.

While this study supports the potential of spinal-level GPR37 activation as a novel therapeutic approach to pain treatment, the following limitations are acknowledged. First, to thoroughly assess the abuse liability of GPR37 agonists, more rigorous testing, such as the intravenous self-administration test, may be necessary, given that commonly abused drugs are administered via routes beyond the intended clinical route. Second, the sensitization of the nociceptive system that maintains chronic pain may differ from that induced by acute injury in its molecular, cellular, and circuit-level mechanisms. Therefore, to preclinically evaluate the therapeutic potential of spinal-level GPR37 agonism for chronic pain treatment, it is necessary to study the effectiveness of i.th. TX14A and PD1 in animal models of chronic pain in future studies.

In conclusion, this study demonstrates that spinal-level GPR37 agonism resolves long-lasting pain hypersensitivity following acute injury by normalizing long-term changes in the spinal nociceptive system (i.e., erasing nociceptive system sensitization), while leaving normal nociception intact. The resolution of acute injury-induced pain hypersensitivity is dependent on GPR37 in TRPV1-lineage DRG neurons. Furthermore, centrally activating this receptor does not show apparent abuse liability in the conditioned place preference paradigm, collectively suggesting that spinally targeting GPR37 in TRPV1-lineage DRG neurons holds promise as a novel pain therapy.

## Disclosures

The authors have no conflicts of interest to declare.

This study was supported by DOD CDMRP under award number W81XWH2110752 (JL) and the NIH HEAL Initiative under award number R61NS127286 (JL).

The data supporting the findings of this study are available upon request.

## Author contributions

**Regan Hammond:** Investigation, Formal analysis, Writing-Original draft. Visualization. **Jigong Wang:** Investigation, Formal analysis. **Ramesh Pariyar:** Investigation, Formal analysis. **Koo Ho:** Resources. **Jun-Ho La:** Conceptualization, Writing-Reviewing and Editing, Supervision, Funding acquisition.

