## Supplemental Figures for "Spinal-level activation of GPR37 in TRPV1-expressing sensory neurons erases nociceptive system sensitization in murine models"

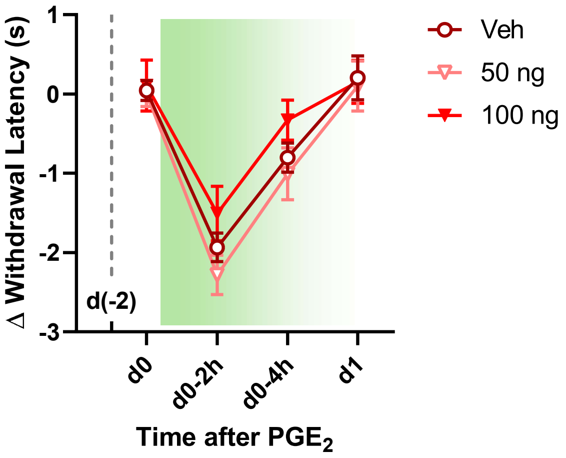


**Suppl Fig 1** (related to Fig 6E). When the individual baseline values were subtracted from the dataset, PD1 at 100 ng, intrathecally administered (gray broken line) 2 days prior to $PGE_2$, had no significant effect on $PGE_2$-induced heat hypersensitivity in naïve (i.e., not-primed) female mice.

**Suppl Fig 1**

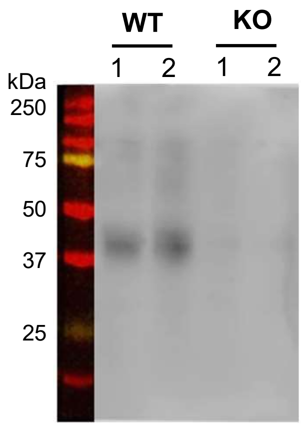


**Suppl Fig 2** (related to Figs 7 and 8). Western blot confirmed the absence of GPR37 in the spinal cord harvested from GPR37 global KO mice.

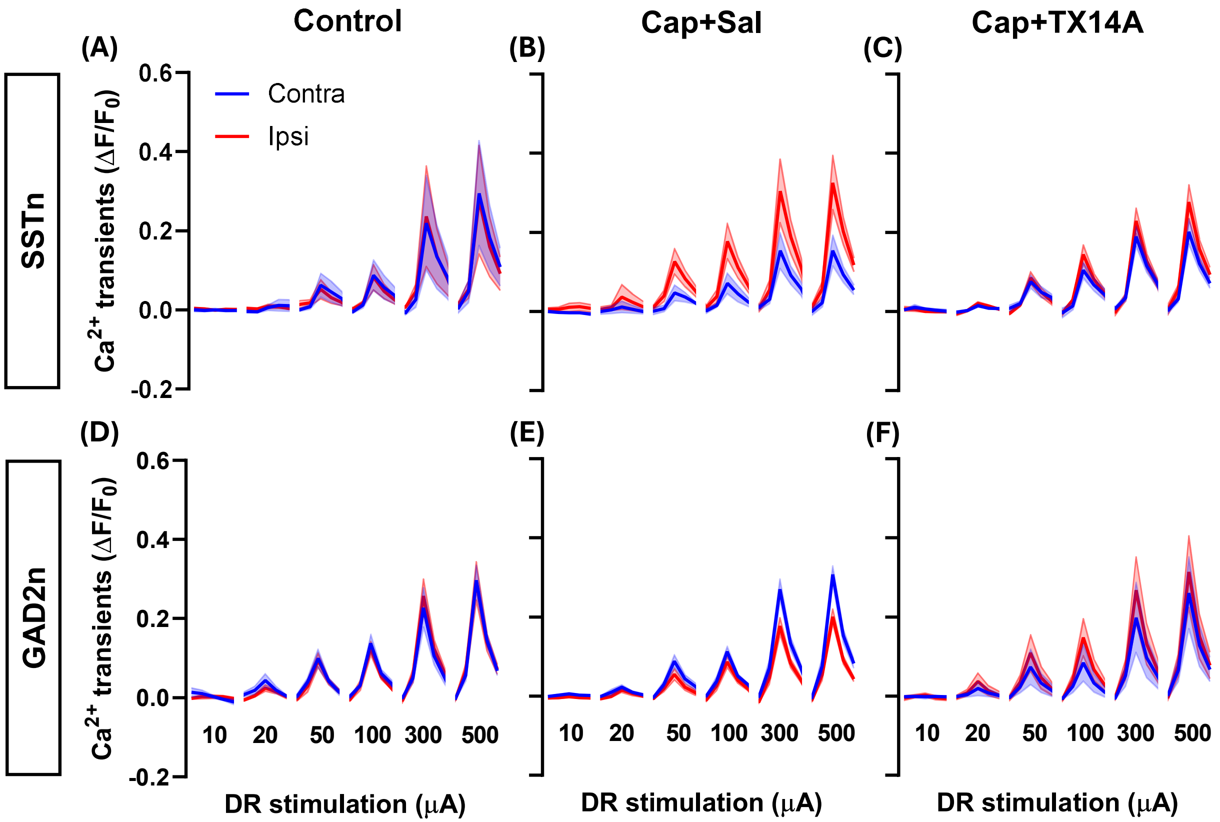


**Suppl Fig 3** (related to Fig 12). In SSTn, there was a trend toward increased $Ca^{2+}$ transients in the ipsilateral dorsal horn of the Cap+Sal group ($F(1,6)=4.943$, $p=0.068$); a significant interaction effect (dorsal horn side x DR stimulation intensity; $F(29,174)=2.864$, $p<0.0001$) was also detected by 2-way RM ANOVA. In GAD2n, $Ca^{2+}$ transients in the Cap+Sal group were significantly decreased in the ipsilateral dorsal horn ($F(1,8)=7.448$, $p=0.026$); a significant interaction effect (dorsal horn side x DR stimulation intensity; $F(29,232)=4.079$, $p<0.0001$) was also detected by 2-way RM ANOVA. For both SSTn or GAD2n in other groups, there was no significant main effect of dorsal horn side or the interaction effect (dorsal horn side x DR stimulation intensity).

**Suppl Fig 3**
